## SUPPLEMENTAL FIGURES for "Host Gastric Corpus Microenvironment Facilitates *Ascaris Suum* Larval Hatching And Infection in a Murine Model"

**Supplement Figure 1**

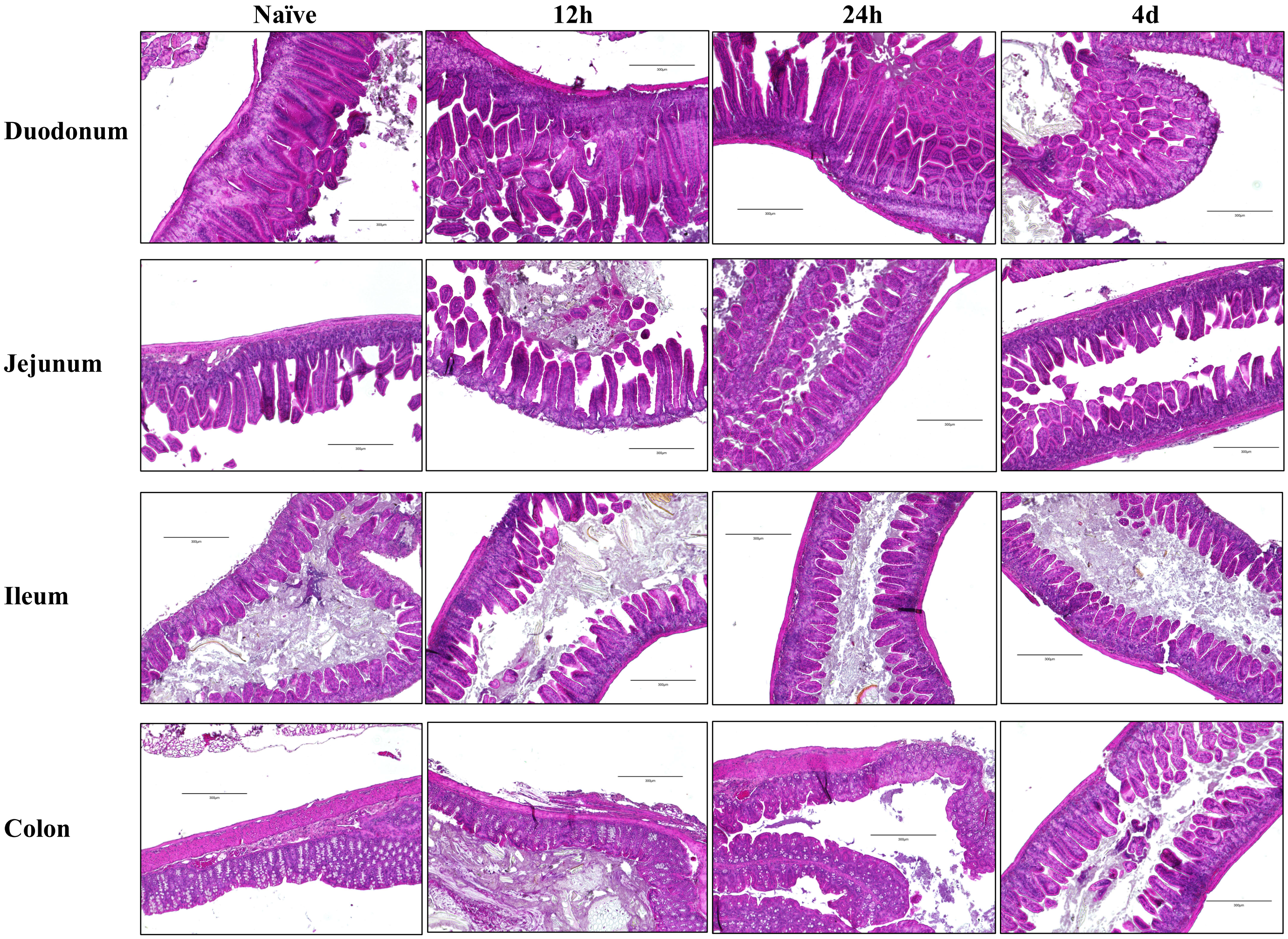


**Supplemental Figure 1: Intestine of mice post *Ascaris* larva infection.** Wildtype mice were oral gavaged with 2,500 *Ascaris* eggs and euthanized at 12 hours, 24 hours or 4 days post infection. H&E staining were carried out on sections of different intestine tissues.

**Supplement Figure 2**

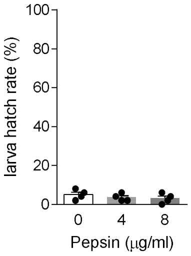


**Supplemental Figure 2: Pepsin does not induce *Ascaris* larva hatching.** *Ascaris* eggs are treated with 4 or 8 μg/ml of pepsin in pH=2 overnight. Larval hatch rate were then calculated.

**Supplement Figure 3**

| | Ctrl | Trypsin | Trypsin+Bile salt | Trypsin+Bile |
|---|---|---|---|---|
| Larval hatch rate % | 0.315789% | 0.27897% | 0.506977% | 0.217391% |

**Supplemental Figure 1: *Ascaris* larva does not hatch in intestinal conditions.** Ascaris eggs are treated in pH=2 for 30 minutes and then in intestinal conditions overnight. Larval hatch rate were then calculated.
